# A pentylenetetrazole-induced kindling zebrafish larval model for evoked recurrent seizures

**DOI:** 10.1101/787580

**Authors:** Sha Sun, Chenyanwen Zhu, Manxiu Ma, Bing Ni, Lin Chen, Hongwei Zhu, Liu Zuxiang

## Abstract

**Background:** Transient pentylenetetrazol (PTZ) treatment on zebrafish larvae has been widely accepted a promising animal model for human epilepsy. However, this model is not ideal due to its acuteness and lack of recurrent seizures, which are the key feature of epilepsy in human disease. It is important to develop a more sensitive zebrafish model for epilepsy with well-controlled, predictable, recurrent seizures.

**New Method:** The new method includes an experimental setup and a treatment protocol. The setup tracks the locomotion activity of up to 48 larvae simultaneously, while a visual stimulus can be presented to each of the 48 animals individually. The protocol treated the larvae through a water bath in 5 mM PTZ while being stimulated with rotating grating stimuli for 1 hour/day from 5 to 7 days postfertilization.

**Results:** The setup captured the locomotion activity of zebrafish larvae during visual stimulation. The new protocol generated recurrent responses after flashing lights 4 hours post PTZ treatment. The effects could be suppressed by the anti-epileptic drug valproic acid. The characteristics of the visual stimulus play a major role in this kindling model.

**Comparisons with Existing Methods:** We compared the proposed method with the transient PTZ model and confirmed that the flashing-light-evoked recurrent seizure is a new feature in addition to the transient changes.

**Conclusions:** The new method generated non-drug-triggered predictable recurrent seizures in response to intermittent photic stimulation in zebrafish larvae and may serve as a sensitive method for anti-epileptic drug screening or a new research protocol in epilepsy research.

## 1. Introduction

Epilepsy, as a common neurological disorder produced by abnormal electrical discharges in the brain, has been characterized by spontaneous recurrent seizures (Stewart et al., 2012). It manifests various types of symptoms, such as cramps in the arms and legs, temporary confusion, or long-lasting convulsions (Galanopoulou et al., 2012). Animal models were applied to address various relevant questions related to the mechanisms of cellular hyperexcitability, synchronous neuronal discharge and rhythmic activity, alterations underlying the transition from an interictal to an ictal state, seizure propagation, seizure cessation, mechanisms and side effects of anti-epileptic drugs (AEDs), and the behavioral disruptions caused by or associated with seizures (Sarkisian, 2001). Unfortunately, approximately one-third of patients suffer from AED-resistant seizures due to different pathway specificities, e.g., Dravet syndrome, even though many new AEDs have been developed based on these animal models over the past 20 years (Griffin et al., 2018; Johan Arief et al., 2018). Epilepsy, as a type of central nervous system disorder, can be only symptomatically treated instead of being cured because a significant number of AEDs target only the symptoms of the disease, and the clear causes of epilepsy during development are poorly understood (Clossen and Reddy, 2017). A hypothesis of the reason for this weakness may be inherited from the model itself, because the same specific mechanisms/pathways are involved during both modeling and screening (Loscher, 2011). Human epilepsy is defined by the appearance of multiple spontaneous recurrent seizures (SRSs), but most animal models are established with seizures induced acutely (Sarkisian, 2001).

Considering the three criteria for the potential ideal animal model for human disease (similar etiology as a human form; the same physiological, behavioral or genetic phenotypes; and the same response to the therapies (Grone and Baraban, 2015), models that generate recurrent seizures are a better choice for understanding epilepsy and AED screening. To achieve that, one approach is using large doses of kainate (a glutamate analog) or pilocarpine (a cholinergic agonist) that induce severe acute seizures followed by the development of spontaneous recurrent seizures several weeks later (Sarkisian, 2001). Another approach is kindling. By repeated application of a stimulus to the limbic system in the amygdala or hippocampus, kindling models induce generalized seizures that are sensitive to the adverse effects of AEDs and are more susceptible to the relevant behavioral and cognitive alterations (Song et al., 2018). Although the kindling model has a special advantage in controlling the seizure focus, depending on the electrode location, its main disadvantage is the extensive kindling stimulation (Samokhina and Samokhin, 2018). In practice, the long-term administration of AEDs and continuous video/electroencephalogram (EEG) monitoring of model animals is technically difficult, expensive, and time-consuming (Loscher, 2011).

Fortunately, zebrafish have been proven to be a rising model for epilepsy (Baraban et al., 2005; Grone and Baraban, 2015; Liu and Baraban, 2019). In the acute seizure zebrafish model, lower concentrations of pentylenetetrazol (PTZ) merely evoked seizure-like behavior, such as increased locomotion activity (Stage I and II) and clonus-like convulsion (Stage III), which can be evoked only in the presence of high concentrations of PTZ (15 mM). Once Stage III is evoked in zebrafish, it cannot be reversed by any method except the administration of AEDs (Baraban et al., 2005). Since this first demonstration of the zebrafish larval model, seizure-like responses have been described by behavioral changes in locomotor activity (Baraban et al., 2005; Barbalho et al., 2016; Berghmans et al., 2007), EEG (Pineda et al., 2011), c-fos expression (Baraban et al., 2005; Barbalho et al., 2016), optical imaging (Turrini et al., 2017), calcium imaging (Liu and Baraban, 2019), and the phosphorylation of ERK1/2 kinases (Randlett et al., 2015). The same PTZ treatment had been proven to be applicable to the adult zebrafish, and the behavioral response profile was characterized by stages that were comparable to those described for the rodent models (Afrikanova et al., 2013; Gupta et al., 2014; Mussulini et al., 2013). Zebrafish have several advantages, such as high homology (70%) to humans (Howe et al., 2013) and being amenable to genetic manipulation (Johan Arief et al., 2018). Moreover, the following features make zebrafish more useful to generate kindling models: they are easy to maintain, they absorb drugs directly from water, and they develop fast (in 3-4 months) (Hortopan et al., 2010). Evidence exists that absence-like seizures are elicited by systematic low-dose PTZ treatments. When the same low dose of PTZ kindling was applied to adult fish, similar cellular responses and cognitive changes were found (Duy et al., 2017; Kundap et al., 2019). However, there is no information on whether the same kindling method could be effective in zebrafish larvae.

In the current study, we aim to develop a zebrafish larva kindling model that is sensitive to intermittent photic stimulation (IPS). IPS is a standard clinical technique during routine diagnostic EEG for determining photosensitivity, especially for some careers such as pilots or traffic controllers (Kasteleijn-Nolst Trenite et al., 2012; Kasteleijn-Nolst Trenite, 2005). Though the prevalence of photoconvulsive seizures related to IPS in the general population is as low as approximately 1 per 10,000 and 1 per 4,000 in individuals between the ages of 5 and 24 years, people known to have epilepsy have from a 2% to 14% chance of having seizures precipitated by light or patterns (Fisher et al., 2005; Martins da Silva and Leal, 2017; Padmanaban et al., 2019; Stewart et al., 2012), with the highest prevalence of photosensitivity reported in juvenile myoclonic epilepsy (JME) at 30.5% (Poleon and Szaflarski, 2017). This means that approximately 1.25% of individuals referred for EEGs might have an abnormal EEG response to certain types of light stimulation (Fisher et al., 2005). Animal models for photosensitive epilepsy have been limited to the baboon (*Papio papio*) (B P-P) and Fayoumi photosensitive chickens (FPCs) for many years, but the precise mechanisms that underscore photosensitivity remain unclear (Poleon and Szaflarski, 2017) and may not closely approximate the human disorder (Fisher et al., 2005). As a special kind of reflex seizure, the photosensitive trait and its underlying mechanisms may enrich our understanding of the transition of normal physiological function to paroxysmal epileptic activity in general (Koepp et al., 2016).

Evidence exists that visual aura is positively correlated with the photoparoxysmal response (PPR), indicating a degree of involvement of the visual cortex during IPS (Martins da Silva and Leal, 2017). It is important to note that up to two-thirds of individuals with photosensitivity are also pattern sensitive (Koepp et al., 2016). The center-surround inhibition in the visual system enables a certain visual stimulus to activate a large number of neurons with high-amplitude neuronal synchronicity (Poleon and Szaflarski, 2017). It consists of the idea that the reflex seizure relies on two important factors: the appropriate afferent volleys and a critical mass of the cortex being activated. More than that, drifting gratings to the center of gaze were not found to be provocative, while gratings that repeatedly change their direction of movement or phase could be highly provocative (Fisher et al., 2005). In combination with the fact that seizure activity associated with cortical dysplasia is often resistant to commonly used AEDs, suggesting that a neuronal migration disorder may also entangle pharmacological mechanisms (Loscher, 2011), it is of special interest to take the impact of visual patterns on kindling into consideration. It is even important for the general population because IPS may evoke seizure-like responses in computer games or video commercials (Fisher et al., 2005). Since the impact of the increased exposure to potentially seizure-triggering visual stimuli may take effect in a more indirect or cumulative way, it will be very useful to have a validated animal model as a screening tool for visual stimuli (Kasteleijn-Nolst Trenite, 2005). Here, we suggest a new kindling method for zebrafish larvae that combines low-dose PTZ with visual pattern stimulation. After three days of kindling, the fish exhibited increased locomotor activity after IPS at 30 Hz.

## 2. Materials and methods

### 2.1. Fish husbandry

Adult zebrafish (*Danio rerio*, AB wild type strain from China Zebrafish Resource Center) were maintained at 28□ under a 14/10 day/night cycle with an automatic fish housing system (ESEN, Beijing, China). All embryos and larvae were raised in E3 embryo medium (egg water is made from instant ocean 60× E3: 17.2 g NaCl, 0.76 g KCl, 2.9 g CaCl_2_.2H2O, 4.9 g MgSO_4_.7H2O dissolved in 1 L Milli-Q water; diluted to 1× in 9 L Milli-Q water plus 100 μl 0.02% methylene blue). Larvae between 5 and 7 days postfertilization (dpf) were used in this study. All animal experimental procedures followed the guidelines of the Institutional Animal Care and Use Committee at the Institute of Biophysics of the Chinese Academy of Sciences (Beijing, China).

### 2.2. Apparatus

#### 2.2.1. Hardware

The setup was designed with standard optic parts and customized 3D-printing components (Fig. 1A). Videos of fish larvae in a 48-well cell culture plate were recorded by IR-sensitive CMOS to track the locomotion activities of the fish. The plate was placed on a customized 3D-printing holder (SI_1) on top of a projector. The projector was mounted to a vertical adjustable stage that allows the visual stimuli to be presented to the projection screen at the bottom of the 48-well plate. A shortpass filter was placed in front of the projector to eliminate the interference of visual stimulus light to the IR CMOS camera while acquiring the activity of fish. Infrared light from surveillance light sources (850 nm) was reflected by a mirror to penetrate through the projection screen and the cell plate from beneath. This design enabled reliable images of fish larvae to be captured by the CMOS camera, while the visual stimulus projected by the projector was visible to the fish but not to the camera. All the parts/components used in this design are summarized in a list for reference (SI_2).

**Fig. 1.**
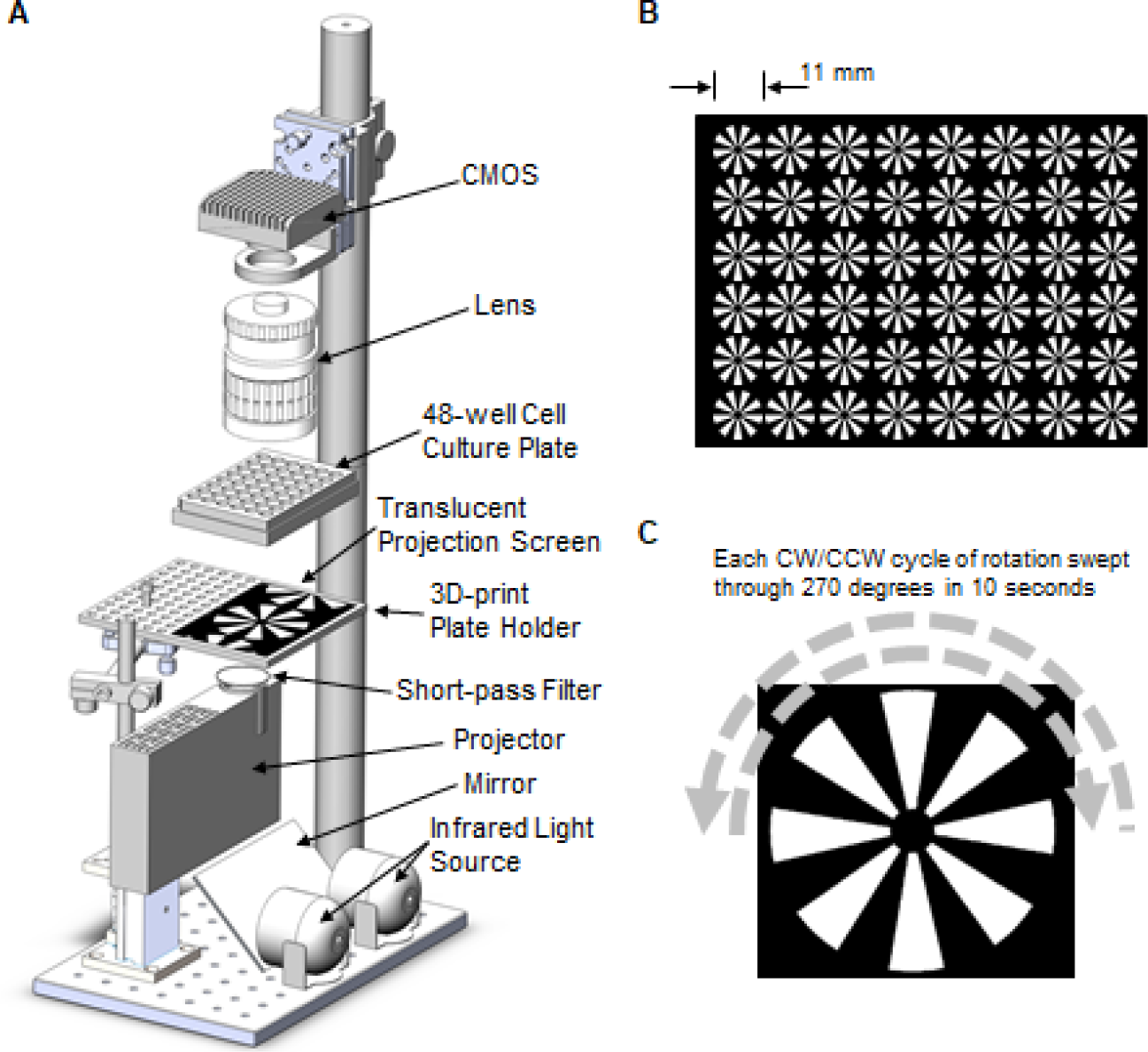
Setup and visual stimulus. (A) With standard optical components and a customized 3D-printed plate holder, the experimental setup enables the tracking of zebrafish larvae in a standard 48-well cell culture plate while visual stimuli were presented from the bottom of the plate. (B) A radial grating was presented to each well individually. The diameter of the grating is equal to the size of the wells in a 48-well plate. (C) The radial grating was rotated with constant velocity and changed direction every 5 seconds.

#### 2.2.2. Visual stimulus

The visual stimuli were generated by MATLAB (MATLAB 2011a, MathWork) and Psychtoolbox-3 (PTB-3). In the proposed protocol, there were 48 rotating gratings arranged in an 8 × 6 array on a black background. The diameter of each grating was 11 mm to fit the cell culture plate (Fig. 1B). The spatial frequency of the grating was 1/45 cycle per degree (Fig. 1C). Each grating rotated clockwise/counterclockwise continuously with a fixed angular velocity that covered 270 degrees in 5 seconds and changed direction (http://www.peterscarfe.com/ptbtutorials.html).

In the flash test, the visual stimulus started with a white blank screen for 20 seconds, followed by 5 flashing sessions. In each session, the screen flashed between black and white at a given frequency for 5 seconds and stayed a white blank screen for 20 seconds. The flashing frequencies for the 5 sessions were 5, 10, 15, 20, and 30 Hz. The total duration of the test was 145 seconds (Fig. 3A).

In the moving dot condition, arenas 11 mm in diameter were presented to each well. There were 4 moving dots in each arena during the 1-hour treatment. The size of the dot was 0.0825 mm. The small dots traveling in line paths covered at least half of the diameter of the arena with a speed of 1.65 mm/s. The onset and paths of the dots were generated randomly in such a way that, though the offset of the dots was independent from one another, a new dot appeared right after the offset of an old one to ensure that there were 4 dots visible in the arena for every moment (Fig. 6A).

### 2.3. Procedure

#### 2.3.1. Dose dependency of PTZ-induced locomotion activity

Fresh working stock (50 mM) was prepared before each experiment by dissolving 0.345 g powdered PTZ (Sigma) in 50 ml E3. For the 1 mM PTZ condition, each well contained 980 μl egg water before 20 μl PTZ working stock was added into the well. For the 5 mM and 15 mM conditions, the combinations of E3 and PTZ working stock were 900 μl + 100 μl and 700 μl + 300 μl, respectively.

For each dose condition, before the PTZ working stock was added into the well, the 48-well plate was recorded with a resolution of 4000◊3000 at a frequency of 2 Hz for 10 minutes. Another recording session with the same parameters was applied after PTZ was added.

After the 10-minute PTZ treatment, a flash test was applied with PTZ still in the well. For the flash test, the IR images were recorded with a resolution of 4000◊3000 at a frequency of 10 Hz. Then, the fish were transferred to a new plate with fresh E3 at 1 ml/well and stayed for 4 hours before another flash test was administered, without PTZ.

#### 2.3.2. PTZ plus visual stimulus protocol

To induce recurrent light-sensitive epilepsy, we designed the PTZ plus visual stimulus protocol (Fig. 4A). The fish were treated with 5 mM PTZ for 1 hour, during which rotating gratings were presented continuously. The treatments were applied from 10 AM to 11 AM from 5 dpf to 7 dpf. After the 1-hour treatment, the fish were transferred to a new plate with fresh E3. A flash test was recorded 4 hours after the last session of PTZ treatment (7 dpf). This protocol will be referred to Grating + PTZ in the following text (Fig. 4B).

To test the recurrent effect of the new protocol, a control condition was used. In the control condition, the timeline was the same as in the Grating + PTZ condition, except that PTZ was applied only for the 1-hour treatments. This control condition will be referred to as PTZ (Fig. 4B).

To test if the other visual stimulus could generate the same effect, a moving dot condition was also applied. In this condition, the parameters were the same as in the Grating + PTZ condition, except that the visual stimuli were moving dots as described above. This condition will be called Dots + PTZ (Fig. 6B).

#### 2.3.3. Anti-epileptic drug treatment

The anti-epileptic drug valproic acid (VPA) was used to verify whether the effects generated by the new protocol were involved in the mechanisms of seizure. Fresh working stock (10 mM) was prepared before the experiments by dissolving 0.083 g powdered VPA (Sigma) in 50 ml E3.

The VPA experiments were based on the Grating + PTZ, PTZ and Dots + PTZ conditions. The timeline and parameters in the VPA experiments were the same as in the previously mentioned experiments, except that a 1 mM concentration was used by adding 100 μl VPA working stock in 900 μl E3 to each well before the flash test (Fig. 5A).

### 2.4. Data analysis

#### 2.4.1. Tracking fish activity in 48-well plates

The IR video clips were analyzed by custom written MATLAB scripts (SI_3). The raw images were first converted into grayscale pictures of 256 gray levels and averaged across each recording session (10 min at 2 Hz or 145 s at 10 Hz) to obtain a background image. Then, a map of standard deviation was generated for each session. For each frame of a recording session, a difference image was generated by subtracting the background from the current frame. Pixels in the difference image with absolute values that were 3 times larger than their corresponding values in the standard deviation map were detected. Those pixels were set to a value of 1, while others were set to a value of 0 before a spatial filter with a Gaussian kernel (size = 5, sigma = 3) was applied. Then, the filtered images were applied to 48 round masks defined on the background image while each mask covered only one well of plate. Pixels with values larger than 0.5 in the mask were detected, and the averaged center of the pixels was defined as the position of the fish relative to the center of this mask/well. The velocity of locomotion activity was calculated by the position differences between two consecutive frames.

#### 2.4.2. Locomotion index in flash tests

In the flash test, the whole experiment was composed of 1 baseline block (the first 20 s), 5 flash blocks (5 s each) and 5 blank blocks (20 s each, Fig. 3A). The mean velocity of locomotion activity was calculated for each block. A global mean of the 11 blocks was used as a threshold to exclude empty wells (some fish failed to survive through the 3-day protocol) or fish that seldom moved (the number of animals in each flash test is emphasized in the figures). The locomotion index (LMi) for each block was defined as a relative value to the global mean.

To test if the locomotion index is different for the 11 blocks, a one-way ANOVA was used. When there was a significant main effect found, an ad hoc test was applied between the baseline block and the other 10 blocks. In spite of that, the responses to the flash itself and the responses during the blank screen after flash were compared by a pairwise t-test for each flashing frequency.

#### 2.4.3. Correlation analysis

For the Grating + PTZ, PTZ and Dots + PTZ experiments, Pearson’s correlations were calculated between the locomotion index of response after a 30 Hz flash, the averaged response for all 5 flash blocks and the averaged response for the other 4 after-flash blocks (Fig. 6).

## 3. Results

### 3.1. PTZ-induced locomotion activity

The PTZ bathing changed the pattern of the locomotion of zebrafish larvae in a concentration-dependent manner (Fig. 2). High concentrations (5 and 15 mM) of PTZ evoked clonus-like convulsive swimming (Stage III, Baraban 2005), while the low concentration (1 mM) did not show an apparent influence on locomotion activity. The onset latency of the changes was shorter at 15 mM than at 5 mM (Fig. 2, lower panel).

**Fig. 2.**
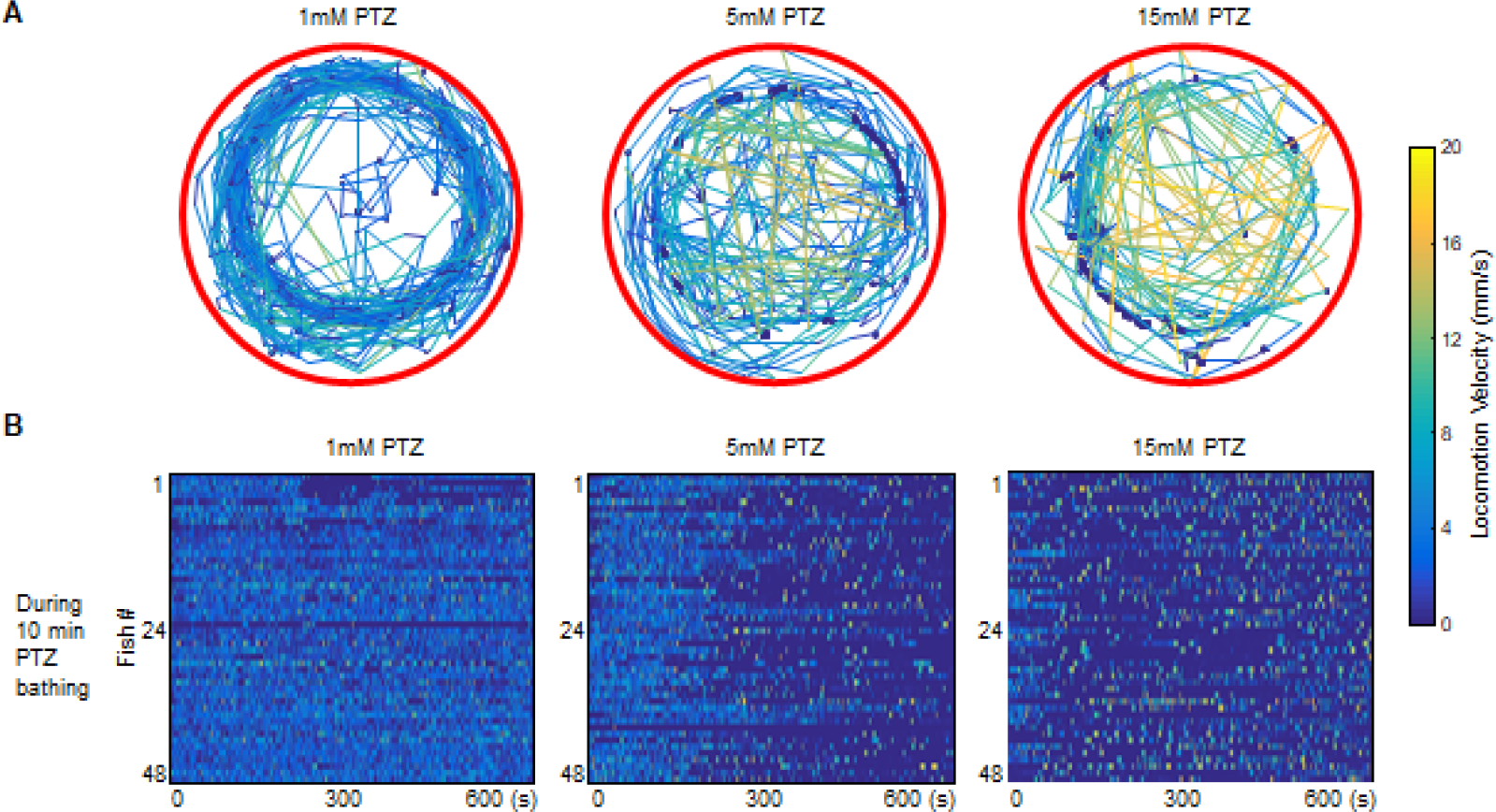
Changes in locomotion activity induced by PTZ are dose dependent. (A) Sample locomotion tracking plots are shown for individual zebrafish in three PTZ concentrations. The color of the line codes locomotion velocity. The number of occasions of the rapid convulsive seizure activity, indicated by yellow lines, increases with PTZ concentration. Plots were obtained from recordings 10 min in duration. (B) PTZ bathing for 10 min changed the locomotion pattern of zebrafish larvae to interleaved clonus-like convulsive swimming with high velocity (Stage III). Notice that the onset of Stage III is dose dependent. Each line in the images represents the activities from one fish.

### 3.2. Light sensitivity measured with flash tests

After a 10 min water bath with PTZ, the zebrafish larvae showed significant responses to flashing light with the presence of PTZ (Fig. 3A). The locomotion index (LMi) was calculated to justify the effects of flashing light. At a concentration of 1 mM, there was a significant main effect of visual conditions (*F*_10,517_ = 3.16, *p* < 0.001), while the response to 15 Hz flashing was higher than baseline (*p* < 0.05). At a concentration of 5 mM, the main effect was significant (*F*_10,506_ = 55.91, *p* < 0.001), and the responses to most flashing frequencies (5, 10, 15, and 20 Hz) were higher than baseline (*p* < 0.001). At a concentration of 15 mM, the main effect was also significant (*F*_10,517_ = 13.79, *p* < 0.001), and the responses to most flashing frequencies were higher than baseline (5 Hz, *p* < 0.05; 10 Hz, *p* < 0.001; 15 Hz, *p* < 0.01; 20 Hz, *p* < 0.001, Fig. 3B, upper panels).

**Fig. 3.**
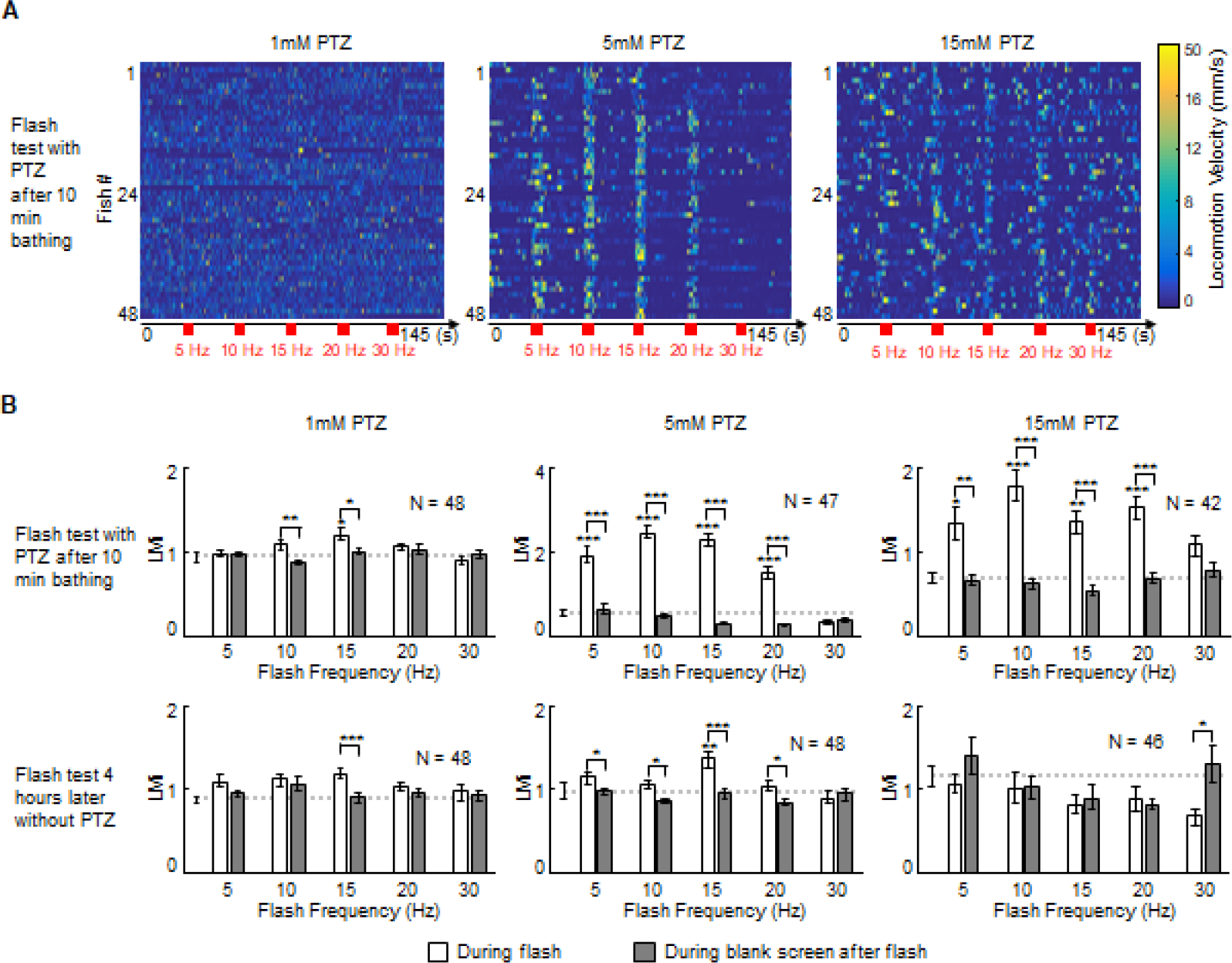
Transient PTZ treatment does not generate recurrent light sensitivity. (A) Light sensitivity tested with a visual flash test. The flash test paradigm is illustrated in each panel. The 5 mM PTZ group showed the strong light sensitivity. (B) The locomotion activity is characterized as the locomotion index (LMi). Upper panels, results of the flash test with PTZ right after the 10 min bathing. The zebrafish larvae showed increased locomotion to the flash itself, but the response remained normal after the flash. Lower panels, results of the flash test 4 hours later without PTZ. Only the 5 mM group showed increased activity in the flash. Dashed line, baseline activity before the first flash. Stars above each bar indicate significant differences from the baseline activity. Stars bridging over two bars indicate significant differences between the responses during the flash and the interval after the flash. Error bars indicate SEM. * p < 0.05, ** p < 0.01, *** p < 0.001.

However, when the same flash test was applied 4 hours later without the presence of PTZ, the response to flashing lights diminished in the 1 mM- and 15 mM-treated fish. Only the treatment of 5 mM PTZ showed significant recurrent responses (*F*_10,517_ = 4.55, *p* < 0.001) to the flashing light of 15 Hz (*p* < 0.01, Fig. 3B, lower panels).

Although none of the transient PTZ treatments elicited recurrent changes in the blank intervals after flashing light, the treatment of 5 mM PTZ induced significant behavioral changes during flashing light immediately after treatment with PTZ and 4 hours post PTZ treatment, which makes the 5 mM treatment a possible candidate to generate recurrent effects (Duy et al., 2017; Kundap et al., 2019).

### 3.3. PTZ plus visual stimulus kindling generated light-sensitive recurrent effects

For the flashing test 4 hours post PTZ treatment at day 7, a significant main effect of visual conditions (*F*_10,418_ = 6.82, *p* < 0.001) was found if the fish were treated with the new PTZ + Grating protocol. Most importantly, the LMi during the interval after 30 Hz flashing was significantly higher than baseline (*p* < 0.01, Fig. 4B, left panel). As a comparison, the fish treated for three days with only PTZ (PTZ condition) showed a different response profile to the flashing light. There was a significant main effect of visual conditions (*F*_10,517_ = 25.90, *p* < 0.001), while the responses to 10, 15, and 20 Hz were significantly suppressed (*p* < 0.001 for 10 and 15 Hz in both flashing and blank intervals, *p* < 0.01 for flashing at 20 Hz). The higher response at 5 Hz was found to be limited to the flashing interval. However, there was no significantly higher response to the blank interval at any of the tested frequencies (Fig. 4B, right panel).

**Fig. 4.**
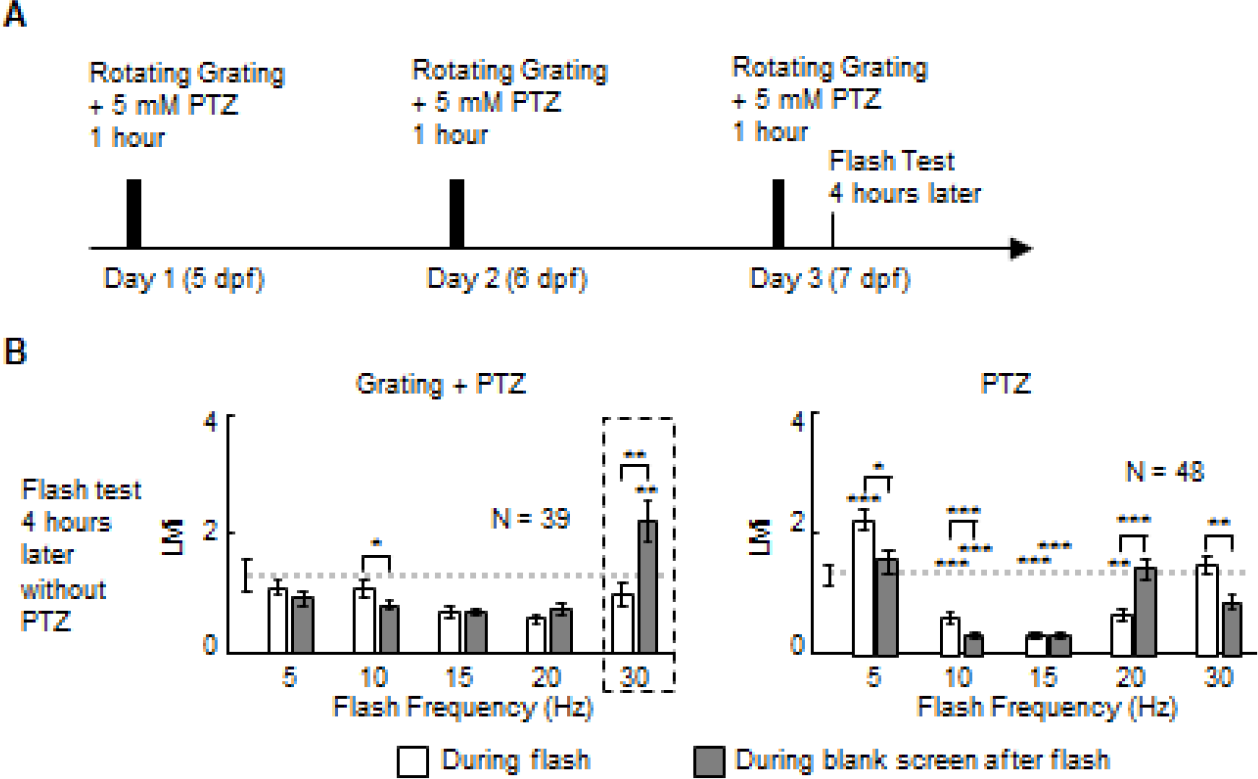
New protocol that generates recurrent light sensitivity. (A) Timeline of the new method. Zebrafish larvae were treated with 5 mM PTZ with visual stimuli for 1 hour every day from 5 dpf to 7 dpf. Flash tests were taken at 4 hours after the last PTZ bathing. (B) The new method induced significantly larger locomotion activity for a time interval after a 30 Hz flash (left panel) for 4 hours. As a comparison, PTZ treatments without visual stimuli for 3 days decreased LMi for several frequency bands (right panel). Error bars indicate SEM. * p < 0.05, ** p < 0.01, *** p < 0.001.

### 3.4. Anti-epileptic drug suppressed light-sensitive recurrent effects

To test whether the recurrent seizure-like behaviors induced by the new protocol can be suppressed by the anti-epileptic drug, we added valproic acid (VPA) to the wells before the flashing test. For the flashing test 4 hours post PTZ treatment at day 7, the application of VPA successfully suppressed the recurrent activity to flashing light. No significant effect was found in the PTZ + Grating + VPA condition (Fig. 5B, left panel). The control condition showed an accumulated suppression of locomotion activity by VPA on the responses found in the PTZ-only condition, with a significant main effect of visual conditions (*F*_10,374_ = 3.54, *p* < 0.001), while the responses to 10, 15, 20, and 30 Hz were significantly suppressed (Fig. 5B, right panel).

**Fig. 5.**
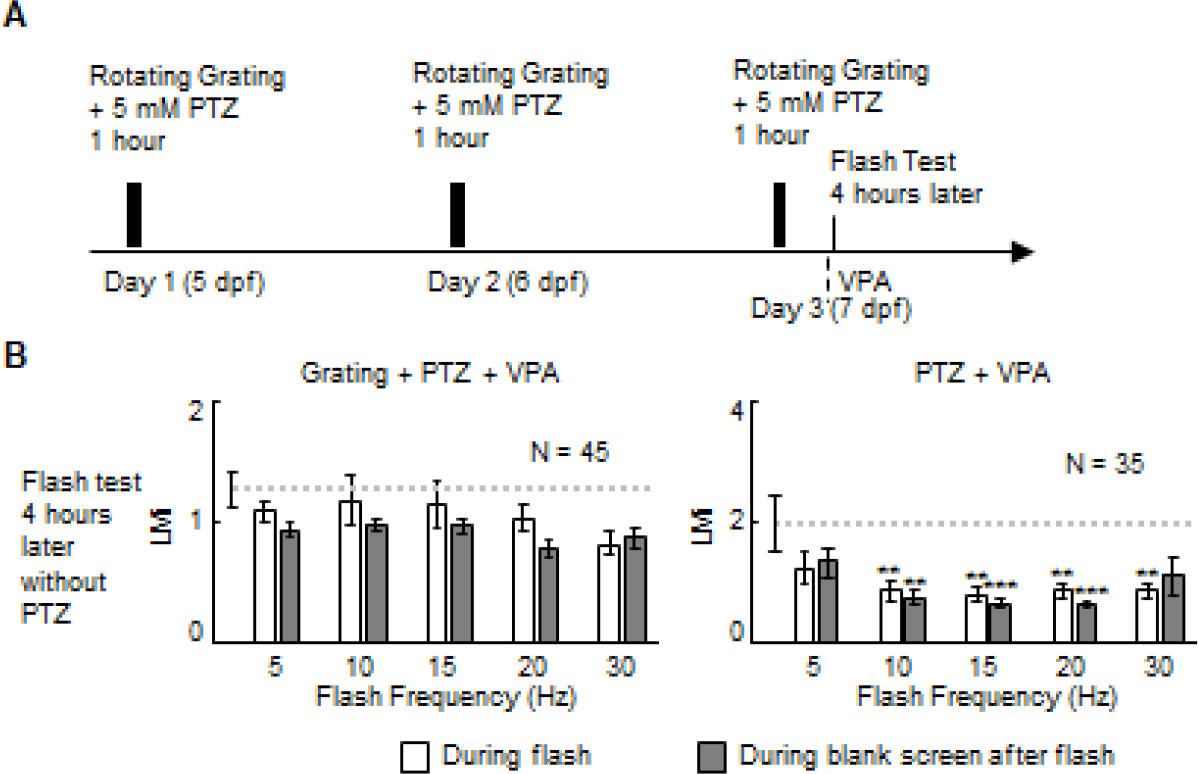
Anti-epileptic drug (VPA) suppressed recurrent light sensitivity. (A) Timeline of the experiment. The conditions were the same as those of the new protocol, except that 1 mM VPA was added to the well before the first flash test and remained in it until the end of the second flash test. (B) The anti-epileptic drug suppressed the recurrent light sensitivity (left panel). As a comparison, 1 mM VPA suppressed the effects of PTZ (right panel). Error bars indicate SEM. * p < 0.05, ** p < 0.01, *** p < 0.001.

### 3.5. Effects of different visual stimuli during kindling

To evaluate whether the features of the visual stimulation of the new protocol play a role in the kindling effect, a different type of visual stimulus was applied as another control condition (Dots + PTZ). In this control condition, the moving dots shared the motion components of the proposed new protocol, but the stimuli remained in a constant direction (Fig. 6A). For the flashing test 4 hours post PTZ treatment at day 7, a significant main effect of visual conditions (*F*_10,429_ = 13.32, *p* < 0.001) was found. The LMi of all the flashing frequencies during or after the flashing was significantly smaller than that of the baseline (*p* < 0.001), except for the interval after 30 Hz flashing (*p* = 0.84, Fig. 6B, left panel). When the anti-epileptic drug VPA was added to the water before the flashing test, the responses to neither the flash nor the interval after flashing were different from baseline (Fig. 4B, right panel), even though a significant main effect was still visible (*F*_10,407_ = 2.48, *p* < 0.01).

**Fig. 6.**
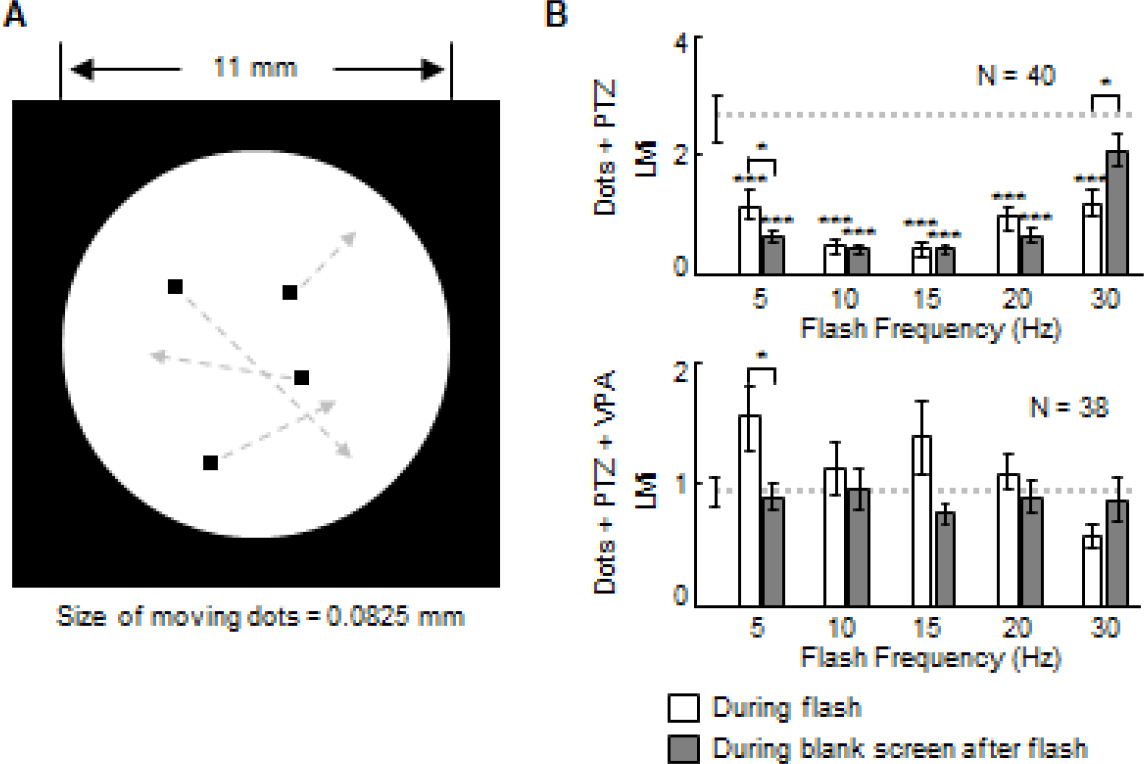
Moving dots with PTZ generate weaker effect. (A) Parameters of the moving dots. For each well, the arena is 11 mm in diameter, in which small dots traveling in line paths covered at least half of the length of the diameter. The onset and paths were generated randomly, but there were 4 moving dots for every moment during the 1 hour treatment. (B) The moving dots with PTZ induced significantly smaller responses to most flash conditions, but the response after the 30 Hz flash was not different from baseline (upper panel). The anti-epileptic drug VPA suppressed the effects induced by the moving dots and PTZ (lower panel). Error bars indicate SEM. * p < 0.05, ** p < 0.01, *** p < 0.001.

### 3.6. Correlation analysis

A significant negative correlation was found between the responses to the interval after 30 Hz flashing and the responses to all flashing intervals (Fig. 7A left panel, *r* = −0.61, *p* < 0.001) or the responses during the interval after flashing for all other frequencies (Fig. 7A middle panel, *r* = −0.67, *p* < 0.001) in the proposed new protocol. The responses to all flashing intervals and the responses during the interval after flashing for all other frequencies showed a significant positive relation (Fig. 7A right panel, *r* = 0.39, *p* < 0.05).

**Fig. 7.**
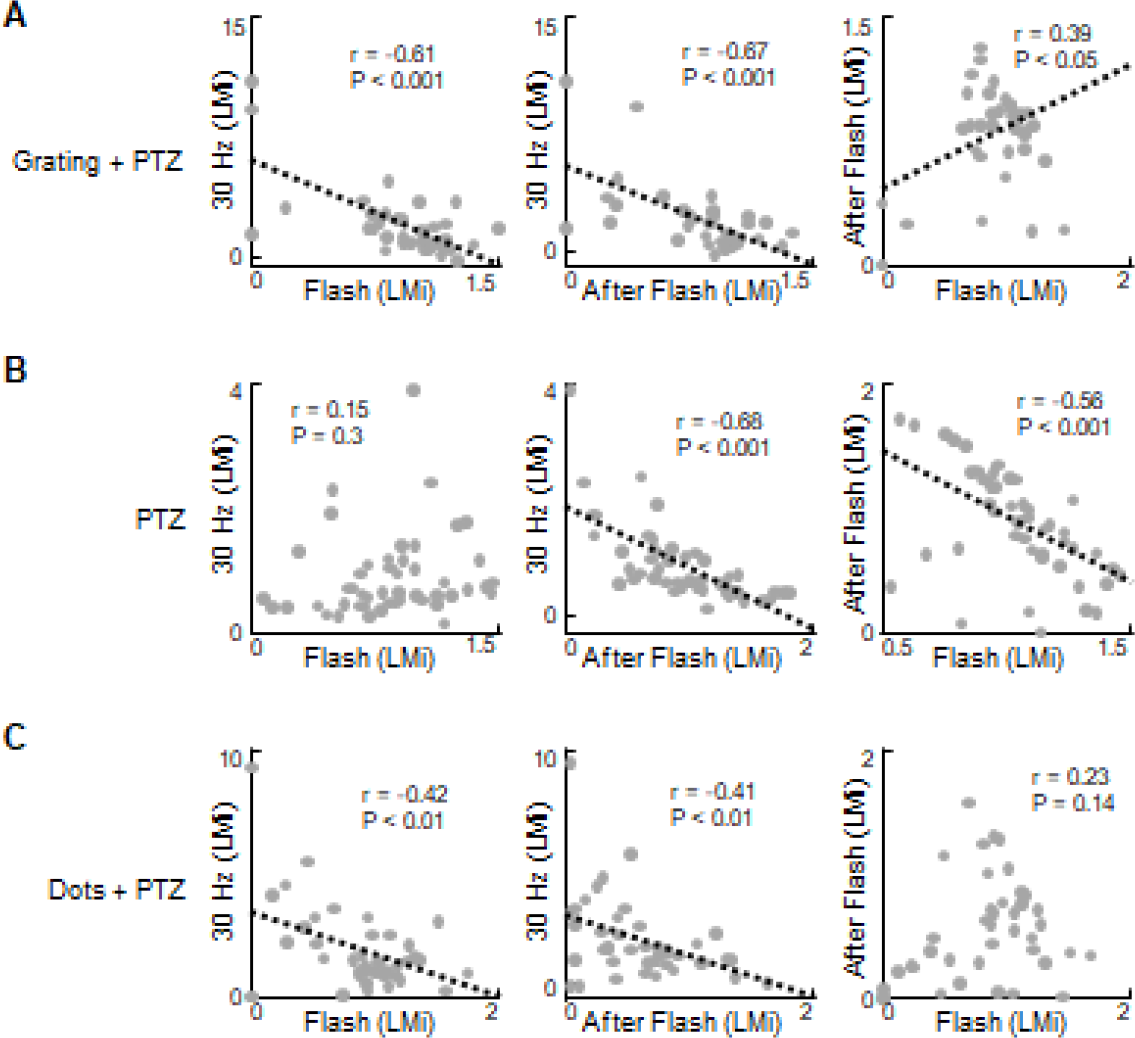
Pairwise correlations between responses to the flash itself and responses after the flash. Left panels, the correlations between responses after the 30 Hz flash and the averaged responses to all flash conditions; middle panels, the correlations between responses after the 30 Hz flash and the averaged responses after the other four frequency bands; right panels, the correlations between the averaged responses to all flash conditions and the responses after the other four frequency bands. The black dashed line shows the linear fit of the dataset if the correlation is significant. Each dot represents the data from one fish.

For the fish treated in the PTZ control condition, although there was a significant negative correlation between the responses to the interval after 30 Hz flashing and the responses during the interval after flashing for all other frequencies (Fig. 7B middle panel, *r* = −0.68, *p* < 0.001), the relation between the responses to all flashing intervals and the responses during the interval after flashing for all other frequencies was also significantly negative (Fig. 7B right panel, *r* = −0.56, *p* < 0.001), which is different from the proposed Grating + PTZ condition.

The visual stimuli control condition (Dots + PTZ) showed similar relations for the first two calculations (Fig. 7C left and middle panels, *r* = −0.42 and *r* = −0.41, *p* < 0.01, respectively), as found in the Grating + PTZ condition, but did not show a significant positive relation in the third calculation (Fig. 7C right panel).

## 4. Discussion

In this paper, we propose a new zebrafish model for epilepsy. By combining PTZ and visual stimulation for three days from 5 dpf to 7 dpf, the new protocol now can induce recurrent seizures after a 4 hour delay by light flashing without the presence of PTZ. Fish showed a significantly higher response to the blank screen after the 30 Hz flashing light. This effect could be eliminated by the anti-epileptic drug VPA. We also demonstrated that the features of the visual stimulus also play a role in this new model.

We developed this new model because existing models have limitations. There are three criteria for the ideal model for epilepsy: similar etiology as a human form; the same physiological, behavioral or genetic phenotypes; and the same response to the therapies. However, existing epilepsy models, though simple and easy to use, failed to fulfill these criteria (Grone and Baraban, 2015). The same problem happened in the zebrafish model for epilepsy. Since the establishment of the PTZ-induced seizure model for epilepsy in zebrafish (Baraban et al., 2005), there have been rich applications (Afrikanova et al., 2013; Banote et al., 2013; Barbalho et al., 2016; Berghmans et al., 2007; Gupta et al., 2014; Hunyadi et al., 2017; Lin et al., 2018; Liu and Baraban, 2019; Mei et al., 2013; Mussulini et al., 2013; Pineda et al., 2011; Siebel et al., 2013; Tiedeken and Ramsdell, 2007; Turrini et al., 2017; Yang et al., 2017). However, the seizures induced by PTZ only cause transient acute seizures, which have poor face validity as the defining feature of epilepsy is spontaneous recurrent seizures (Johan Arief et al., 2018). To address this issue, PTZ kindling has been applied to adult zebrafish. Similar to the kindling model of mice (Song et al., 2018), animals begin having spontaneous recurrent seizures and continue to have prolonged seizures (Duy et al., 2017; Kundap et al., 2019). One labor-consuming but useful feature of PTZ kindling is that the process takes several weeks, which makes it possible to study the subtle mechanisms of epileptogenesis, including the chemokine receptor ligand family (Liu et al., 2007), cellular responses (Duy et al., 2017), morphological changes of neural networks (Samokhina and Samokhin, 2018) and cognitive functions (Kundap et al., 2019). In the current paper, we extend the zebrafish kindling model from the adult to the larva, which is facilitated by the fast development of the larva and its visual sensitivity to light.

The visual startle response of zebrafish larva, characterized by rapid body movement responses to sudden decreases in brightness, indicates a kind of light sensitivity (Emran et al., 2008). However, in the acute PTZ model of zebrafish larvae, light stimulation has always been applied as a condition lasting for several minutes, in the context of dark incubation before and/or after it (Lin et al., 2018), and the level of locomotion activity might depend on illumination conditions (Yang et al., 2017). In contrast to this long-lasting illumination condition, IPS is a more generally used stimulus for human tests (Kasteleijn-Nolst Trenite, 2005), and the evoked responses have been categorized as photomyoclonic responses (PMRs, stimulus-locked) and photoconvulsive responses (PCRs, continuing discharges may evolve to overt epileptic seizures), each of which may involve different neural networks (Martins da Silva and Leal, 2017). We believe that for a successful kindling model of zebrafish larvae, it is necessary to take the evoked responses of IPS into consideration because animal models for photosensitivity and epilepsy have been limited to the baboon (*Papio papio*) (B P-P) and Fayoumi photosensitive chickens (FPCs) for many years (Martins da Silva and Leal, 2017).

There are other reasons for developing a light-sensitive model of epilepsy. The underlying mechanisms of light-sensitive seizures may be related to the transition from normal physiological function to paroxysmal epileptic activity, since PPR has been reported to occur in 0.5-8.9% of healthy individuals (Koepp et al., 2016). It also consists of the idea that photosensitivity might be found in all epilepsy types, suggesting adaptive clinical trial strategies and a faster/less-expensive screening of new AEDs (Padmanaban et al., 2019). Considering the fact that valproic acid (VPA) and levetiracetam (LEV) have been used in the treatment of photosensitivity with varying degrees of success (Poleon and Szaflarski, 2017), even though AEDs are not suggested for photosensitive patients in general, it is promising to develop more finely grained methods based on light sensitivity to detect less than ideal anti-epileptic compounds that have the potential to be effective AEDs after chemical modification (Johan Arief et al., 2018). Moreover, since IPS-provoked clinical symptoms and motor events are always of short duration, it is possible to test the same animal many times with different intervals without the influence of PTZ itself.

It is beyond the scope of this paper to explore the potential mechanisms underlying this new model. There are two factors we believe to be important. The first is the involvement of visual brain area because there is a close relationship between visual aura and the photoparoxysmal response (Stewart et al., 2012). In the baboon model, several neuroimaging studies have revealed that some brain regions and cortico-subcortical networks substitute seizures for the IEDs induced by IPS (Martins da Silva and Leal, 2017). DTI and EEG-fMRI indicate microstructural changes in photosensitive patients. The second factor is regional hyperexcitability. Transcranial magnetic stimulation (TMS) studies have indicated that patients with photosensitivity have a regional hyperexcitability of the primary visual cortex and that there was increased BOLD activation within the visual, premotor and parietal cortices before the onset of PPR after IPS (Padmanaban et al., 2019). It is assumed that the normal brain needs strong external interventions to reach the seizure state, but the functional or structural reorganization of underlying neuronal networks in pathological conditions can move normal activities closer to a lowered seizure threshold (Koepp et al., 2016). It is straightforward to speculate that our PTZ plus visual stimulus paradigm introduced regional hyperexcitability in the visual area of zebrafish larvae. The sensitivity to 30 Hz flashing lights in the PTZ plus rotating grating condition, in comparison with the PTZ-only condition, suggested a developmental hyperexcitability of the visual area. Regions of cortical hyperexcitability receiving appropriate afferent volleys and a critical mass of the cortex being activated are two preconditions in these cortical triggered reflex seizures (Ferlazzo et al., 2005). Since the moving dot condition in our experiments showed similar but weaker effects than the rotating grating condition, while the flashing lights as triggering events showed the same effects for both conditions, the difference between them may rely on weaker hyperexcitability induced by moving dots during the three days of PTZ kindling. This difference, from another perspective, confirmed the regional hyperexcitability in the visual area.

In the future, it is possible to evaluate these factors from a perspective of system dynamics to explore a wide array of possible biophysical mechanisms for seizure genesis (Jirsa et al., 2014), such as neuron/pathway stress levels, neurotransmitter differences, etc. As demonstrated in the correlation analysis, the locomotion index of the interval after a 30 Hz flashing light is negatively correlated with either the responses to the intervals during flashing light or the responses to the interval after other frequencies in both the PTZ plus rotating grating and PTZ plus moving dot conditions. As a comparison, the PTZ-only condition showed a different correlation pattern. Since the responses to the flashing light itself and the continuing responses after flashing light should be supported by different neural networks, the new model may be underscored by abnormal system dynamics (Liu and Baraban, 2019).

## Conclusion

The treatment of zebrafish with PTZ has become a promising animal model of epilepsy. Kindling may induce recurrent seizures that are similar to human symptoms. Photosensitivity, as a reflexive property evaluated during routine clinical EEG recording, needs to be considered in models of epilepsy. The new model here suggested a combined PTZ and visual stimulus treatment for 3 days of kindling and successfully generated an epilepsy model of recurrent seizures that is sensitive to IPS. Increased locomotion activity after flashing light of 30 Hz demonstrated its potential as a sensitive index to be used in AED screening or to probe the neural networks involved in seizure onset and propagation.

