## Supplementary material for "A pentylenetetrazole-induced kindling zebrafish larval model for evoked recurrent seizures": PartsList.docx

**Supplementary Table 1. Components of the experimental setup for the PTZ+VIS paradigm**

| Part Number | Qty | Description | Parameters/info | Supplier |
| --- | --- | --- | --- | --- |
| GCM-3013M | 1 | Optical Breadboards | 200 mm x 300 mm | Daheng Optics |
| GCM-1602M | 1 | Vertical Manual Stage |  | Daheng Optics |
| GCM-030105M | 1 | Post | Ø12mm, length 203mm | Daheng Optics |
| GCM-1903M | 1 | Manual Yaw Platform |  | Daheng Optics |
|  | 1 | Plate Holder | Customized 3D-printing |  |
|  | 1 | Plate Holder Spacer | Customized 3D-printing |  |
| GCT-080315 | 1 | Translation Lens Mount |  | Daheng Optics |
| GCM-030103M | 1 | Post | Ø12mm, length 102mm | Daheng Optics |
| GCM-550101 | 2 | 90°Post Clamp |  | Daheng Optics |
| GCM-030101M | 2 | Post | Ø12mm, length 51mm | Daheng Optics |
| Coolux X5 | 1 | Projector |  | Coolux Technology |
|  | 1 | M6 nut |  | Daheng Optics |
|  | 1 | M6 bolt | 40mm | Daheng Optics |
| GCM-220503M | 1 | Post Clamp |  | Daheng Optics |
|  | 1 | 48-well plate |  | NEST Biotechnology |
| MV1200 | 1 | CMOS |  | Beijing PDV Instrument |
| LENS-CM1140P-8M | 1 | Zoom Lens |  | Uniview Technologies |
| GCM-221102M | 2 | Post | Ø38.1mm, length 355mm | Daheng Optics |
| FM01R | 1 | Short-pass Filter |  | Thorlabs |
