## Supplementary material for "A pentylenetetrazole-induced kindling zebrafish larval model for evoked recurrent seizures": DataAnalysis01_CalulcateMeanStd.m

```
clear;


basepathname = '';
[filename pathname]=uigetfile({'*.raw','À´×Ô¸ßËÙ¾µÍ·µÄÊý¾Ý';...
    '*.*','ALL FILES(*.*)'},'ÇëÑ¡ÔñÍ¼ÏñÎÄ¼þ',basepathname,'MultiSelect', 'on');


if isequal([filename pathname],[0,0])
    return;
end

if(iscell(filename)==0)
    T=filename;
    clear filename;
    filename{1}=T;
end

filestem=filename{1};
DataFormat = 2;
if(DataFormat==0)
    ResolutionX=320;
    ResolutionY=240;
    filestem=filestem(1:9);
    for i=1:length(filename)
        filename{i}=sprintf('%s%d.raw',filestem,i);
    end
    FileId=1;
    HeaderOffset=0;
    FrameOffset=0;
elseif (DataFormat==1)
    ResolutionX=512;
    ResolutionY=384;
    filestem=filestem(1:14);
    for i=1:length(filename)
        filename{i}=sprintf('%s%d.raw',filestem,i);
    end
    FileId=1;
    HeaderOffset=0;
    FrameOffset=0;
else
    K=strfind(filename{1},'at_');
    filestem=filestem(1:K(1)+2);
    K=regexp(filename{1},'at_(\w*)x(\w*)_F(\w*)_', 'tokens');
    ResolutionX=str2num(K{1}{1});
    ResolutionY=str2num(K{1}{2});
    FileId=str2num(K{1}{3});
    for i=1:length(filename)
        filename{i}=sprintf('%s%dx%d_F%d_%d.raw',filestem,ResolutionX, ResolutionY, FileId, i);
    end
    HeaderOffset=8;
    FrameOffset=12;
end

disp(filename);
FrameNumbers=zeros(length(filename),1);
for i=1:length(filename)
    str=[pathname filename{i}];
    disp(str);
    rawinfo=dir(str);
    FrameNumbers(i)=(rawinfo.bytes-HeaderOffset)/(ResolutionX*ResolutionY+FrameOffset);
end
basepathname=pathname;


im111=zeros(ResolutionX, ResolutionY);
str=[pathname filename{1}];
fid=fopen(str);
fseek(fid,0,'bof');
if(DataFormat==2)
    TimingFreq=fread(fid,1,'*int64');
    TimingTag=fread(fid,1,'*int64');
    ExternalTriggerTag=fread(fid,1,'*int32');
else
    TimingFreq=0;
    TimingTag=0;
    ExternalTriggerTag=0;
end
A=fread(fid, ResolutionX*ResolutionY, '*uint8');
B=reshape(A,ResolutionX,ResolutionY);
%Brgb=demosaic(B','grbg');
Brgb=demosaic(B','bggr'); % 'gbrg' 'grbg' 'bggr' 'rggb'
C=squeeze(Brgb(:,:,1))';
im111 = C;
imgbase = double(im111)/256;
figure;imagesc(imgbase);

%%

ImgMean=zeros(ResolutionX, ResolutionY);

frameCount=0;
for i=1:length(filename)
    str=[pathname filename{i}];
    disp(str);
    fid=fopen(str);
    fseek(fid,0,'eof');
    filelength=ftell(fid);
    if(filelength==-1)
        disp('Error in file length test.');
        disp(filein);
        pause;
        exit;
    end
    for j=1:FrameNumbers(i)
        disp([i j]);
        fseekstatus=fseek(fid, (j+1)*(ResolutionX*ResolutionY+FrameOffset)+HeaderOffset, 'bof');
        if(fseekstatus==0)
            fseekstatus=fseek(fid, j*(ResolutionX*ResolutionY+FrameOffset)+HeaderOffset, 'bof');
            if(DataFormat==2)
                TimingTag=fread(fid,1,'*int64');
                ExternalTriggerTag=fread(fid,1,'*int32');
            else
                TimingTag=0;
                ExternalTriggerTag=0;
            end
            A=fread(fid, ResolutionX*ResolutionY, '*uint8');
            frameCount=frameCount+1;
            B=reshape(A,ResolutionX,ResolutionY);
            Brgb=demosaic(B','bggr');
        %    C=squeeze(mean(Brgb,3));
            C=squeeze(Brgb(:,:,1))';
            C=double(C);
            ImgMean=ImgMean+C;
        end
    end
    fclose(fid);
end
ImgMean=ImgMean/frameCount;

%%
ImgStd=ImgMean;
ImgStd(:)=0;
frameCount=0;
for i=1:length(filename)
    str=[pathname filename{i}];
    disp(str);
    fid=fopen(str);
    fseek(fid,0,'eof');
    filelength=ftell(fid);
    if(filelength==-1)
        disp('Error in file length test.');
        disp(filein);
        pause;
        exit;
    end
    for j=1:FrameNumbers(i)
        disp([i j]);
        fseekstatus=fseek(fid, (j+1)*(ResolutionX*ResolutionY+FrameOffset)+HeaderOffset, 'bof');
        if(fseekstatus==0)
            fseekstatus=fseek(fid, j*(ResolutionX*ResolutionY+FrameOffset)+HeaderOffset, 'bof');
            if(DataFormat==2)
                TimingTag=fread(fid,1,'*int64');
                ExternalTriggerTag=fread(fid,1,'*int32');
            else
                TimingTag=0;
                ExternalTriggerTag=0;
            end
            A=fread(fid, ResolutionX*ResolutionY, '*uint8');
            frameCount=frameCount+1;
            B=reshape(A,ResolutionX,ResolutionY);
            Brgb=demosaic(B','bggr');
        %    C=squeeze(mean(Brgb,3));
            C=squeeze(Brgb(:,:,1))';
            C=double(C);
            D=C-ImgMean;
            ImgStd=ImgStd+D.*D;
        end
    end
    fclose(fid);
end
ImgStd=ImgStd/frameCount;
ImgStd=sqrt(ImgStd);
%%
figure;
subplot(2,2,1);imagesc(ImgMean);
subplot(2,2,2);imagesc(ImgStd);
colormap('gray');

filenameout=[pathname 'MeanAndStd.mat'];
save(filenameout,'ImgMean','ImgStd');
```
