## Supplementary material for "A pentylenetetrazole-induced kindling zebrafish larval model for evoked recurrent seizures": FlashTest.m

```
% Clear the workspace
close all;
clear all;
tic;

SerialPort=serial('COM2','BaudRate',9600);

try
    fopen(SerialPort);
catch err
    fprintf('COM2´ò¿ªÊ§°Ü¡£\n');
end

if(strcmp(SerialPort.Status,'open'))
    for si=1:5
        fwrite(SerialPort,0);
    end
end


AssertOpenGL;
Screen('Preference', 'SkipSyncTests', 1);
Screen('Preference', 'ConserveVRAM', 64);
HideCursor;
% Here we call some default settings for setting up Psychtoolbox

FlashDurationSecs = 5;
RestDurationSecs = 20;

% Get the screen numbers
screens = Screen('Screens');

% Draw to the external screen if avaliable
screenNumber = max(screens);

% Define black and white
white = WhiteIndex(screenNumber);
black = BlackIndex(screenNumber);
grey = white / 2;
inc = white - grey;

% Open an on screen window
%[window, windowRect] = PsychImaging('OpenWindow', screenNumber, grey);
[window, windowRect] = Screen('OpenWindow',screenNumber, black);
% [window, windowRect] = Screen('OpenWindow',screenNumber, black,[0 0 1024 768]);

% Get the size of the on screen window
[screenXpixels, screenYpixels] = Screen('WindowSize', window);


background=zeros(screenYpixels, screenXpixels);
background(:)=black;
checkers=zeros(screenYpixels, screenXpixels);
checkers(:)=white;

% Make the checkerboard into a texure (4 x 4 pixels)
checkerTexture = Screen('MakeTexture', window, checkers);
backgroundTexture = Screen('MakeTexture', window, background);


% Sync us and get a time stamp
vbl = Screen('Flip', window);
waitframes = 0.5;
% Hz: 5 --> 12 frames
%    10 -->  6
%    15 -->  4
%    20 -->  3
%    30 -->  2

FlashFreqSettings=[11 5 3 2 1];

% Maximum priority level
%topPriorityLevel = MaxPriority(window);
%Priority(topPriorityLevel);
ifi = Screen('GetFlipInterval', window);
Screen('DrawTextures', window, backgroundTexture);
vbl = Screen('Flip', window);
%pause;

Screen('DrawTextures', window, checkerTexture);
if(strcmp(SerialPort.Status,'open'))
    for si=1:5
       fwrite(SerialPort,1);
    end
end
vbl = Screen('Flip', window);
startTime = vbl;
while vbl - startTime < RestDurationSecs
    Screen('DrawTextures', window, checkerTexture);
        % Flip to the screen
    vbl = Screen('Flip', window);
end

t0=toc;
for n=1:length(FlashFreqSettings)
    if(strcmp(SerialPort.Status,'open'))
        for si=1:5
            fwrite(SerialPort,10+n);
        end
    end
    vbl = Screen('Flip', window);
    startTime = vbl;
    while vbl - startTime < FlashDurationSecs
        Screen('DrawTextures', window, backgroundTexture);
        % Flip to the screen
        vbl  = Screen('Flip', window, vbl + (waitframes - 0.5) * ifi);
        Screen('DrawTextures', window, checkerTexture);
        % Flip to the screen
        vbl  = Screen('Flip', window, vbl + (FlashFreqSettings(n) - 0.5) * ifi);
    end
    if(strcmp(SerialPort.Status,'open'))
        for si=1:5
            fwrite(SerialPort,30+n);
        end
    end
    startTime = vbl;
    while vbl - startTime < RestDurationSecs
        Screen('DrawTextures', window, checkerTexture);
        % Flip to the screen
        vbl = Screen('Flip', window);
    end
    disp(toc);
end

Screen('DrawTextures', window, backgroundTexture);
Screen('Flip', window);
%pause;
Screen('CloseAll');

if(strcmp(SerialPort.Status,'open'))
    fclose(SerialPort);
end


return;
```
