## Supplementary material for "A pentylenetetrazole-induced kindling zebrafish larval model for evoked recurrent seizures": MultiMovingDotTest48Cells.m

```
% Clear the workspace
close all;
clear all;
tic;

Parameter=load('MultiMovingDotTest48Cells.txt');


AssertOpenGL;
Screen('Preference', 'SkipSyncTests', 1);
Screen('Preference', 'ConserveVRAM', 64);
HideCursor;
% Here we call some default settings for setting up Psychtoolbox

%PsychDefaultSetup(1);
movieDurationSecs = 5;
restDurationSecs = 2;
T = 5; % Stop-to-stop interval in sec
A = 180; % Stop-to-stop swept angle in deg
peakVelocity = A/pi;

% Get the screen numbers
screens = Screen('Screens');

% Draw to the external screen if avaliable
screenNumber = max(screens);

% Define black and white
white = WhiteIndex(screenNumber);
black = BlackIndex(screenNumber);
grey = white / 2;
inc = white - grey;

% Open an on screen window
%[window, windowRect] = PsychImaging('OpenWindow', screenNumber, grey);
[window, windowRect] = Screen('OpenWindow',screenNumber, black);
% [window, windowRect] = Screen('OpenWindow',screenNumber, black,[0 0 1024 768]);

% Get the size of the on screen window
[screenXpixels, screenYpixels] = Screen('WindowSize', window);


% Get the centre coordinate of the window
xCenter=floor(screenXpixels/2);
yCenter=floor(screenYpixels/2);

% Set up alpha-blending for smooth (anti-aliased) lines
Screen('BlendFunction', window, 'GL_SRC_ALPHA', 'GL_ONE_MINUS_SRC_ALPHA');
% Query the frame duration
ifi = Screen('GetFlipInterval', window);
% Screen resolution in Y
screenYpix = windowRect(4);
% Number of white/black circle pairs
rcycles = 0.5;
% Number of white/black angular segment pairs (%integer%)
tcycles = 8;
% Define a simple 4 by 4 checker board
xylim = 2 * pi * rcycles;
[x, y] = meshgrid(-xylim: 2 * xylim / (screenYpix - 1): xylim,...
    -xylim: 2 * xylim / (screenYpix - 1): xylim);
at = atan2(y, x);
checks = ((1 + sign(sin(at * tcycles) + eps)...
    .* sign(sin(sqrt(x.^2 + y.^2)))) / 2) * (white- black) + black;

checks(:)=white;

circle = x.^2 + y.^2 <= xylim^2;
circle2 = x.^2 + y.^2 <= xylim^2/40;
circle = circle .* ~circle2;
checks = circle .* checks + black * ~circle;


% Make the checkerboard into a texure (4 x 4 pixels)
checkerTexture = Screen('MakeTexture', window, checks);


% We are going to draw four textures to show how a black and white texture
% can be color modulated upon drawing
yPos = yCenter;
xPos = xCenter; %linspace(screenXpixels * 0.2, screenXpixels * 0.8, 4);

% Define the destination rectangles for our spiral textures. Each scaled by
% 50x
[s1, s2] = size(checks);
baseRect = [0 0 s1 s2] .*1;
dstRects = nan(4, 48);
%dstRects(:, 1) = CenterRectOnPointd(baseRect, xPos, yPos);
dstRects(1,1)=baseRect(1)+floor(xPos/2);
dstRects(2,1)=baseRect(2)+floor(yPos/2);
dstRects(3,1)=baseRect(3)+floor(xPos/2);
dstRects(4,1)=baseRect(4)+floor(yPos/2);

%MaskOffsetX=100;
%MaskOffsetY=25;
%MaskGridSizeX=196;
%MaskGridSizeY=170;
%MaskSize=100;

MaskOffsetX=Parameter(1);
MaskOffsetY=Parameter(2);
MaskGridSizeX=Parameter(3);
MaskGridSizeY=Parameter(4);
MaskSize=Parameter(5);

DotSize=Parameter(6);
DotSpeed=Parameter(7);
DotNumber=Parameter(8);


for gridi=1:8
    for gridj=1:6
        dstRects(1,(gridi-1)*6+gridj)=MaskOffsetX+(gridi-1)*MaskGridSizeX+1;
        dstRects(2,(gridi-1)*6+gridj)=MaskOffsetY+(gridj-1)*MaskGridSizeY+1;
        dstRects(3,(gridi-1)*6+gridj)=MaskOffsetX+(gridi-1)*MaskGridSizeX+MaskSize;
        dstRects(4,(gridi-1)*6+gridj)=MaskOffsetY+(gridj-1)*MaskGridSizeY+MaskSize;
    end
end

DotInfo=zeros(DotNumber,9); %startx starty endx endy stepcount offsetx offsety currentx currenty

% Switch filter mode to simple nearest neighbour
filterMode = 0;

% Set the inital rotation angle randomly and in increment per frame to 3
% degrees
angles = rand(1, 5) .* 360;

% Sync us and get a time stamp
vbl = Screen('Flip', window);
waitframes = 0.5;

% Maximum priority level
%topPriorityLevel = MaxPriority(window);
%Priority(topPriorityLevel);
ifi = Screen('GetFlipInterval', window);
%pause;
t0=toc;
A=zeros(1000,1);
A(1:2:999)=1;
A(2:2:1000)=-1;
for n=1:1000
    vbl = Screen('Flip', window);
    degPerFrame = A(n);
    startTime = vbl;
    x = 0;
    % Batch Draw all of the texures to screen
    while vbl - startTime < movieDurationSecs
        t = vbl - startTime;
        
        for dotid=1:DotNumber
            if(DotInfo(dotid,5)==0)
                posA=rand(10,2);
                posB=rand(10,2);
                disAB=zeros(100,1);
                for Aid=1:10
                    for Bid=1:10
                        disAB((Aid-1)*10+Bid)=sqrt((posA(Aid,1)-posB(Bid,1))^2+(posA(Aid,2)-posB(Bid,2))^2);
                    end
                end
                tmpMask=disAB;
                tmpMask(:)=1;
                tmp=find(disAB<0.5);
                tmpMask(tmp)=0;
                tmp=find(disAB>1.2);
                tmpMask(tmp)=0;
                disAB=disAB.*tmpMask;
                tmp=find(disAB>0);
                I=tmp(1);
                tmpDis=disAB(I);
%                [tmpDis,I]=max(disAB);
                posA=posA*MaskSize;
                posB=posB*MaskSize;
                tmpDis=tmpDis*MaskSize;
                DotInfo(dotid,1)=posA(floor((I-1)/10)+1,1);
                DotInfo(dotid,2)=posA(floor((I-1)/10)+1,2);
                DotInfo(dotid,3)=posB(mod((I-1),10)+1,1);
                DotInfo(dotid,4)=posB(mod((I-1),10)+1,2);
                DotInfo(dotid,5)=floor(tmpDis/DotSpeed);
                DotInfo(dotid,6)=(DotInfo(dotid,3)-DotInfo(dotid,1))/DotInfo(dotid,5);
                DotInfo(dotid,7)=(DotInfo(dotid,4)-DotInfo(dotid,2))/DotInfo(dotid,5);
                DotInfo(dotid,8)=DotInfo(dotid,3);
                DotInfo(dotid,9)=DotInfo(dotid,4);
            else
                DotInfo(dotid,8)=DotInfo(dotid,1)+DotInfo(dotid,5)*DotInfo(dotid,6);
                DotInfo(dotid,9)=DotInfo(dotid,2)+DotInfo(dotid,5)*DotInfo(dotid,7);
                DotInfo(dotid,5)=DotInfo(dotid,5)-1;
            end
        end
        
        checks2 = zeros(MaskSize,MaskSize);
        checks2(:) = white;
        for dotid=1:DotNumber
            tmpx1=floor(DotInfo(dotid,8)-DotSize/2);
            tmpy1=floor(DotInfo(dotid,9)-DotSize/2);
            tmpx2=tmpx1+DotSize;
            tmpy2=tmpy1+DotSize;
            if(tmpx1<1)
                tmpx1=1;
            end
            if(tmpx1>MaskSize)
                tmpx1=MaskSize;
            end
            if(tmpy1<1)
                tmpy1=1;
            end
            if(tmpy1>MaskSize)
                tmpy1=MaskSize;
            end
            if(tmpx2<1)
                tmpx2=1;
            end
            if(tmpx2>MaskSize)
                tmpx2=MaskSize;
            end
            if(tmpy2<1)
                tmpy2=1;
            end
            if(tmpy2>MaskSize)
                tmpy2=MaskSize;
            end
            checks2(tmpx1:tmpx2,tmpy1:tmpy2)=black;
        end
        
        Screen('Close',checkerTexture);
        checkerTexture = Screen('MakeTexture', window, checks2);
        % Batch draw our scaled, rotated and color modulated textures
        Screen('DrawTextures', window, checkerTexture, [], dstRects);
        % Flip to the screen
        vbl  = Screen('Flip', window, vbl + (waitframes - 0.5) * ifi);
    end
    while vbl - startTime < restDurationSecs
            vbl = Screen('Flip', window);
    end
    disp(toc);
end

Screen('Flip', window);
%pause;
Screen('CloseAll');
return;
```
